# Ontology pre-training improves machine learning-based predictions for metabolites

**DOI:** 10.1101/2025.09.30.679573

**Authors:** Charlotte Tumescheit, Martin Glauer, Simon Flügel, Martin Larralde, Fabian Neuhaus, Till Mossakowski, Janna Hastings

## Abstract

Recent advances in the field of machine learning have shown that integration of expert knowledge improves performances, in particular for complex domains such as biology. Bio-ontologies offer a rich source of curated biological knowledge that can be harnessed to this end. Here, we describe an intuitive and generalisable approach to embed the knowledge contained in a classification hierarchy derived from a bio-ontology into a machine learning model as an intermediate training step between general-purpose pre-training and task-specific fine-tuning in a process that we call ‘ontology pre-training’. We show that this approach leads to an improvement in predictive performance and a reduction in training time for a broad range of prediction tasks relevant to understanding metabolite functions in living systems, using a range of datasets derived from MoleculeNet. We see the biggest improvement for regression tasks, e.g. prediction of lipophilicity and aqueous solubility of molecules, and a robust improvement for most classification tasks. Our approach can be adapted for a wide range of knowledge sources, models and prediction tasks.

## Introduction

Breakthrough performance has been achieved for many of the grand challenges in computational biology through recent advances in deep learning, including, most famously, the Nobel Prize-winning AlphaFold system [1, 2]. These important achievements have raised hopes that the technology may significantly accelerate discoveries across a wide range of open biological challenges, including single cell and spatial -omics [3], and RNA biology [4]. However, for many important prediction tasks in biology, only relatively small and noisy datasets are available for training models, smaller than what might be ideal for data-hungry deep learning architectures [3].

Incorporation of prior biological knowledge can augment machine learning-based predictions and reduce the need for huge datasets, substantially improving predictive performance and robustness. Prior knowledge has been shown to improve many different prediction challenges, from predicting protein function [5] to inferring molecular causality [6]. However, while there are many existing approaches to incorporate prior knowledge into neural network-based machine learning systems, these are often task-specific, or require custom architectures [7]. Ontologies are computable representations of entities and their interrelationships [8] that can be used for a wide range of applications in data science and artificial intelligence. In the domain of biology, bio-ontologies encode a wealth of biological knowledge in the form of a hierarchy of labelled, defined, annotated, and often richly interrelated classes [8]. The most widely used among these is the Gene Ontology [9], which encodes the functions and activities of genes in a species-independent way that drives functional interpretation in the molecular life sciences. Other widespread bio-ontologies include the ChEBI ontology of biologically interesting chemistry [10] and the human phenotype ontology [11], which provides a formalisation of distinct phenotypes in a cross-species fashion enabling translational research.

Bio-ontologies are a natural source of prior knowledge for predictive models. However, in order to use this knowledge in machine learning models, they need to be appropriately “embedded” into the vector representation space of such models [12]. Existing methods for embedding ontologies into predictive models typically involve traversal and iterative encoding of the ontology structure, content and labels. A commonly used method is OWL2Vec [13], which embeds the contents of an ontology represented in the Web Ontology Language (OWL, https://www.w3.org/OWL/). However, while ontology embedding methods create an embedded version of the contents of the ontology, they do not necessarily align that knowledge with the underlying data type for which the knowledge from the ontology is relevant to enhance predictions. Alternative approaches are therefore needed to align the embedded representation of the ontology with the quite different embedded representation of other input data types. In protein function prediction for example, predictive models are usually trained solely on protein sequences, which are represented by a protein language model in a latent space structured in terms of sequence features only. The learned embeddings are therefore disjoint of the knowledge encoded in function representation ontologies such as the Gene Ontology. In metabolite property prediction, models trained only on the chemical structure of molecules similarly learn a representational space different from one for the knowledge encoded in metabolite ontologies. Methods for ontology embedding often do not provide a direct means to connect these different input data types [12].

We propose an intuitive approach harnessing some of the knowledge from an ontology to augment predictive models through a dedicated training process that teaches the model the structure of the ontology in direct association with the input data type. We call our approach ‘Ontology pre-training’, and it consists of a training step designed to teach the model the structure of the ontology in association to the input data type by training the model to predict ontology classifications for the input data type. Our approach requires only that the underlying input data type for the predictive model can be associated with the relevant ontology classes, whether through ontology annotations, through metadata embedded in the ontology, or some other form of association. The approach then creates a dedicated training strategy that embeds knowledge from the ontology into a predictive model in a way that is able to support the further specialisation of the model for downstream predictive tasks.

We apply our approach to a case study on the structure-based prediction of the biologically relevant properties of small molecule metabolites, using the ChEBI ontology [14] as the source of relevant knowledge. ChEBI is a large ontology for biologically relevant small molecules, and includes both a structure-based classification and a classification of relevant biological roles. Crucially for our purpose, it contains structural annotations in the form of SMILES strings [15], which at the leaf level of the ontology represent the structures of small molecules. The same structural annotations are commonly used as input data in molecular property prediction models and chemical language models [16].

In a pilot case study on a single dataset previously presented [17], we showed preliminary evidence that this approach was able to improve model predictive performance and reduce training time for the multi-class predictions of toxicity, a challenging molecular prediction task of broad relevance for the biosciences, drug discovery and ecology. In the current work we go beyond this single case study to show that the approach works more broadly, extending our evaluation to a range of prediction problems including additional classification tasks as well as a number of regression tasks, as found in the MoleculeNet dataset [18]. We also evaluate the method on an extended solubility dataset for aqueous solubility prediction, beyond that available in MoleculeNet, as aqueous solubility is one of the key parameters informing bioavailability, yet its accurate prediction remains a difficult challenge [19].

In the next section, we describe our methods and the datasets used in the pre-training as well as the evaluation in detail. Thereafter, we present the results of our experiments, followed by a discussion including a comparison to prior work.

## Methods

### Experimental Design

The overall objective of our experiments is to compare the predictive performance, as well as the fine-tuning training time and convergence, starting from two comparable base models, namely one with and one without ontology pre-training, across a range of different prediction tasks. To achieve this objective, we train an initial base model with self-supervised pre-training, and then create an extended version with ontology pre-training, then apply fine-tuning to both of the base models for each of the tasks. The overall training workflow is illustrated in **Fig. 1**, and the different datasets used and training steps applied are further elaborated below.

**Fig. 1.**
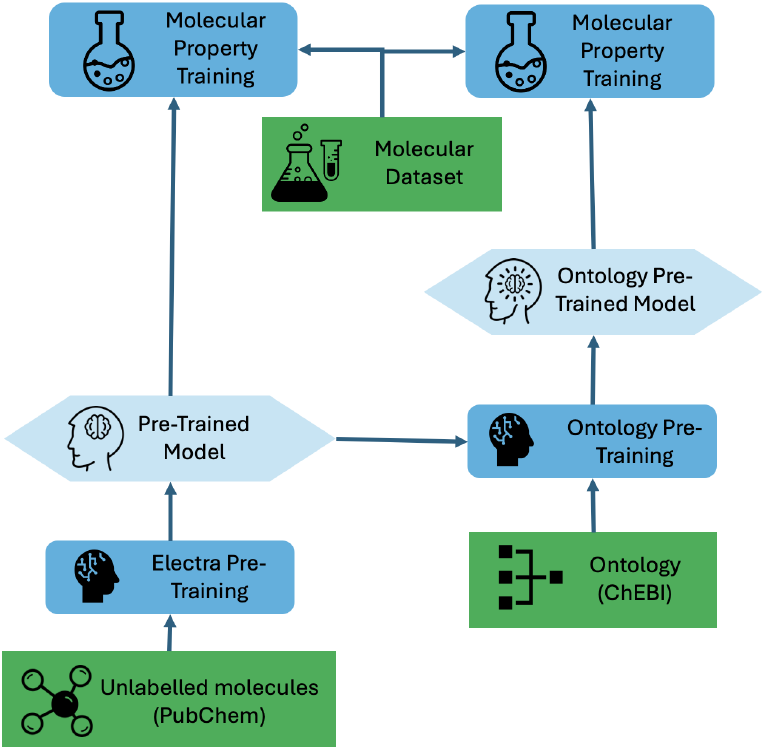
Training stack for the overall experimental design in which a standard pre-trained model is compared to an ontology pre-trained model in subsequent training and prediction tasks.

### Datasets

We are using several different datasets for the different steps in the overall experimental design (**Table 1**).

**Table 1.** Overview of datasets used for training. *a: Randomly subsampled from the full dataset*.

| Dataset | Task | # Labels | Size |
| --- | --- | --- | --- |
| PubChem | Pre-Training | - | 200,000<sup>a</sup> |
| ChEBI | Ontology Pre-Training | 997 | 185,007 |
| Aqueous Solubility | Regression | 1 | 17,339 |
| ESOL | Regression | 1 | 1,128 |
| FreeSolv | Regression | 1 | 642 |
| Lipophilicity | Regression | 1 | 4,200 |
| Tox21 | Classification | 12 | 7,831 |
| ClinTox | Classification | 2 | 1,484 |
| SIDER | Classification | 27 | 1,427 |
| BBBP | Classification | 1 | 2,050 |
| BACE | Classification | 1 | 1,513 |

### PubChem

**PubChem** is a comprehensive open database for chemical information [20], containing more than 100 million distinct chemical records. From this database, we randomly extracted 200, 000 molecular representations, in the form of SMILES strings, for our pre-training. SMILES offers a compact linear representation of the molecular structure of a molecule that is widely used as the input representation format for molecular language models [15]. For example, the SMILES string for the bioactive molecule *caffeine* is ‘Cn1cnc2n(C)c(=O)n(C)c(=O)c12’.

### ChEBI

Chemical Entities of Biological Interest (**ChEBI**) is an ontology containing 61, 181 curated and 123, 826 non-curated compounds (ChEBI release 231) organised in a hierarchy of chemical classes [14]. For the ontology pre-training step, we derived a dataset from ChEBI using the SMILES of the 185, 007 compounds, as well as their flattened classification in the ChEBI hierarchy, resulting in a labelled dataset suitable for multi-label classification, where our model is tasked of predicting the ChEBI class labels from an input SMILES string. To enable satisfactory learning, we retained only classes with sufficient support: we extracted all subclasses of *molecular entity* in ChEBI that had at least 100 members containing SMILES strings, resulting in 185, 007 compounds and 997 classes, analogously to our previous work [17].

### Aqueous solubility - Combining solubility datasets

In [21] seven datasets of thermodynamic and kinetic aqueous solubility data were cleaned and curated according to a custom pipeline and metrics. We inspected all these datasets separately again by using a simple regression and a simple deep neural network (DNN) to explore the performance of a naive model for each dataset. We discarded two datasets that had a comparatively worse performance (based on RMSE for the regression and F1 score for the DNN). We thus retained the following datasets for evaluating our approach on aqueous solubility prediction: **AQSOL** [22], **AQUA** [23, 24], **ESOL** [25], **OCHEM** [26], **PHYS** [27].

### MoleculeNet

**MoleculeNet** [18] is a collection of datasets designed for benchmarking machine learning models for molecular property predictions across a range of tasks. In the current work, we focus on a subset of MoleculeNet that uses only SMILES as the input data type, excluding those that require 3D coordinates or other additional inputs (**Table 1**). Of those tasks, we select both regression tasks and classification tasks.

We use the following regression datasets. **ESOL** [25] consists of water solubility values. **FreeSolv** [28] consists of values of hydration free energy of small molecules in water. The **Lipophilicity** dataset contains lipophilicity values, which inform membrane permeability and solubility, derived from the ChEMBL database [29].

We use the following classification datasets. **Tox21** [30] describes toxicity values on 12 different targets. **SIDER** [31] describes adverse drug reactions for 27 system organ classes. **ClinTox** [32] consists of drug compounds’ toxicity as well as whether they have been approved or failed by the FDA. **BBBP** [33] contains the blood-brain barrier permeability property. **BACE** [34] describes binding results for a set of inhibitors.

While 3 of these datasets are balanced, the remaining 2 datasets feature class imbalance as well as missing labels (**Table 2**).

**Table 2.** Ratio of negative (0) and positive (1) labels as well as missing labels (n/a) for classification datasets.

| Dataset | 0 | 1 | n/a |
| --- | --- | --- | --- |
| Tox21 | 72,084 (76.71%) | 5,862 (6.24%) | 16,026 (17.05%) |
| SIDER | 16,661 (43.24%) | 21,868 (56.76%) | 0 (0.00%) |
| ClinTox | 1,466 (49.39%) | 1,502 (50.61%) | 0 (0.00%) |
| BBBP | 483 (23.56%) | 1567 (76.44%) | 0 (0.00%) |
| BACE | 822 (54.33%) | 691 (45.67%) | 0 (0.00%) |

### Model training

In the last decade, the growth in parameter size of deep-learning models has led to the adoption of a transfer learning methodology which consists of training a model on large amounts of unannotated data, in a step known as *pre-training*. The resulting pre-trained model is then adapted to more specialized tasks in a later step known as *fine-tuning* using much fewer data. For large language models in particular, the pre-training task takes the form of either a generative task, where the model must predict the next token of a sentence (GPT-style models [35]), or a discriminative task, where the model must reconstruct a masked or corrupted input (BERT-style models [36]).

#### Step 1: Standard pre-training

The intention behind the general model pre-training is to provide the model with a distributional understanding of the input data and common patterns that it might encounter. By predicting masked tokens or detecting substitutions, the model learns to understand the context in which tokens likely occur and to reconstruct likely combinations of tokens. This training step does not involve any higher level semantics in the form of categorisation or labelling of the input. In this work, the pre-training allows the model to gain an understanding of the SMILES language [15], which features a special syntax for e.g. rings or functional groups.

We use ELECTRA [37], a Transformer-based [38] model, for initial pre-training, leveraging its replaced token detection objective rather than the standard masked language modeling task. In this architecture, two Transformer models are trained concurrently: a *generator* that performs a masked language task, and a *discriminator* that is trained to detect the substitutions performed by the generator. For this initial task, we used the 200, 000 molecules randomly sampled from the PubChem database [20] described earlier with a random 85/2.25/12.75 training/validation/test split.

#### Step 2: Ontology pre-training

To incorporate relevant knowledge from a bio-ontology into the pre-trained model, we add a second pre-training step. The strategy we use to incorporate the knowledge from the ontology into the base model is to design a hierarchical multi-class classification training objective. The latter is based on the ontology classification as a set of labels for associated input molecules.

For the ontology pre-training step, we use data from the ChEBI ontology [14] described earlier, namely 185, 007 SMILES strings with 997 associated labels each. We use the iterative stratification algorithm developed in [39] to stratify our data according to the labels (i.e., the ChEBI superclasses), ensuring that each label appears similarly often in each split.

The ontology pre-training is implemented in the ChEB-AI^1^ Python library, which also supports pre-processing of chemical data, including datasets extracted from ChEBI [14] and PubChem [20]. In addition, it provides a training environment for a variety of model architectures, including transformers. Notably, the latest version of ChEB-AI features an update to PyTorch 2.0 [40], a rewrite of the framework integrating the PyTorch Lightning library [41] to support distributed training, and an update of the ChEBI dataset from release 200 to release 231.

#### Step 3: Fine-tuning on various molecular prediction tasks

Finally, we fine-tune both models (the model with ontology pre-training as trained in step 1+2, and the model without ontology pre-training as trained with only step 1) to a number of classification and regression tasks to assess the effect of integrating the ontology to downstream tasks. The tasks are: (1) predicting hydration free energy of small molecules in water using the FreeSolv dataset; (2) predicting membrane permeability and solubility using the Lipophilicity dataset; (2)predicting aqueous solubility using the FreeSolv dataset; (3)predicting aqueous solubility using the expanded solubility dataset; (5) predicting binding results for a set of inhibitors using the BACE dataset; (6) predicting blood-brain barrier permeability property using the BBBP dataset; (7) predicting adverse drug reactions using the SIDER dataset; (8) predicting drug’s toxicity and FDA approval using the ClinTox dataset; (9) predicting toxicity using the Tox21 dataset.

As loss functions, we use the mean squared error (MSE) for regression tasks and binary cross-entropy (BCE) for classification tasks, which are the most common loss functions for both task types respectively.

Hyperparameters (**Table 3**) are chosen to be consistent for all training steps, however, to combat over-fitting, which is common when learning from small and noisy datasets, for the last training step we apply a lower learning rate, as well as higher hidden and token dropouts. All data and code used in this study is publicly available^2^.

**Table 3.** Hyperparameters used when training models. *Step 1: Standard pre-training, step 2: ontology pre-training, step 3: fine-tuning*.

| Hyperparameter | Step 1 and 2 | Step 3 |
| --- | --- | --- |
| Vocab Size | 1400 | 1400 |
| Hidden Size | 256 | 256 |
| Number of attention heads | 8 | 8 |
| Number of hidden layers | 6 | 6 |
| Epochs | 100 | 100 |
| Learning rate | $10^{-3}$ | $10^{-4}$ |
| Hidden dropout | 0.1 | 0.4 |
| Token dropout | 0.0 | 0.2 |
| Optimizer | Adamax | Adamax |

### Evaluation and Metrics

To evaluate the effect of adding an ontology pre-training step, we fine-tune two models for a number of datasets: the first model is simply trained on PubChem (Step 1), whereas the second model is trained on PubChem followed by the ontology pre-training (Step 1+2). Then, both these models are fine-tuned in the same way on each dataset (Step 1+2+3 versus Step 1+3). To evaluate the impact of the ontology pre-training, we compare the final predictive performance between these two models as well as the evolution of the training loss over the epochs. We consider the average of three runs for different seeds for each fine-tuning step.

Concerning metrics, for regression tasks, we report the coefficient of determination (*R*^2^) and the Root Mean Square Error (RMSE) values to compare the models. *R*^2^ measures the correlation between the true and predicted values, the best possible score is 1. RMSE measures the the difference between the true and predicted values, the lower the RMSE value the better the performance.

For classification tasks, we report the (micro-/macro-)F1 score and receiver operating characteristic area under the curve (ROC-AUC) score to compare the models. The F1 score is the harmonic mean of precision and recall for a binary classifier; for a multi-label classifier the micro-F1 score treats each label as a prediction from a binary prediction, and computes the F1 score for all predictions at once, while the macro-F1 score averages over the F1 scores computed for each label independently. The receiver operating characteristic curve shows the true positive rate against the false positive rate for each threshold, the best possible ROC-AUC score is 1, while 0.5 means that the classifier is as good as random guesses. The ROC-AUC is usually suitable to evaluate balanced classification problems only[42], which is the case of some datasets, but not all of them (**Table 2**).

To obtain stratified splits without data leakage, we group the SMILES based on their MHFP6 fingerprint [43], which we calculated using RDKit [44], and the Hamming distance between them. In some cases, this delayed over-fitting as we achieved more balanced training/test/validation sets.

## Results

### Regression tasks

For all of the regression task datasets, we observe improved predictive performance in the ontology pre-trained model as compared to the model with only standard pre-training. The predictive results for all regression datasets are reported in **Table 4**. E.g. for Lipophilicity dataset, we observe an improvement in the RMSE from 0.905 without the ontology pre-training to 0.750 with the ontology pre-training step, as well as an improvement in *R*^2^ from 0.403 to 0.591.

**Table 4.** Test set evaluation for regression tasks. *R*^2^ and RMSE metrics, averaged over three runs, with standard deviation. Better model in bold. *OPT: Ontology Pre-Training*.

| Dataset | OPT | RMSE | $R^2$ |
| --- | --- | --- | --- |
| FreeSolv | w/ | **1.367** $\pm 0.080$ | **0.880** $\pm 0.014$ |
| | w/o | 1.570 $\pm 0.091$ | 0.841 $\pm 0.018$ |
| Lipophilicity | w/ | **0.750** $\pm 0.014$ | **0.591** $\pm 0.015$ |
| | w/o | 0.905 $\pm 0.015$ | 0.403 $\pm 0.020$ |
| ESOL | w/ | **0.719** $\pm 0.034$ | **0.863** $\pm 0.013$ |
| | w/o | 0.832 $\pm 0.015$ | 0.817 $\pm 0.007$ |
| Expanded Solubility Dataset | w/ | **0.869** $\pm 0.021$ | **0.859** $\pm 0.007$ |
| | w/o | 0.978 $\pm 0.038$ | 0.821 $\pm 0.014$ |

Furthermore, for each of the datasets, we observe more stable training, i.e. there is less fluctuation of the metrics between consecutive epochs (**Fig. 2** for the Lipophilicity dataset, **Fig. S1-3** for the remaining datasets).

**Fig. 2.**
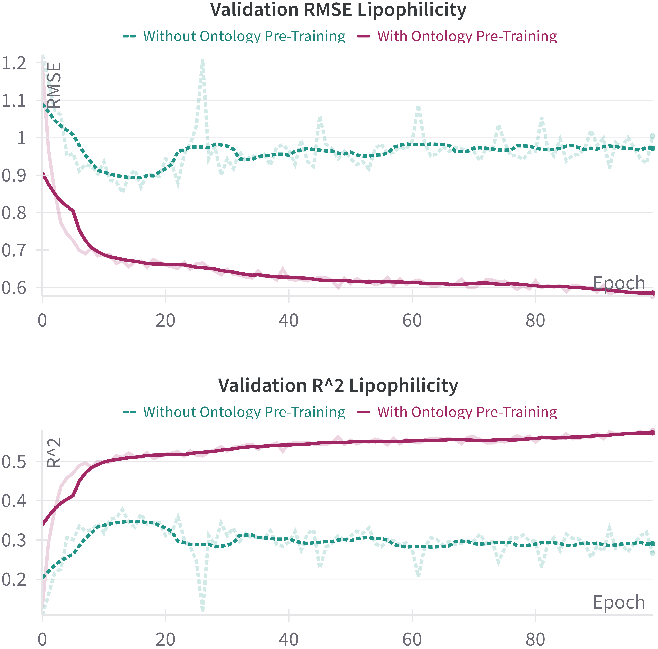
Development of RMSE, *R*^2^ during training per epoch on the validation dataset for Lipophilicity, using a model trained without (turquoise, dashed) and with ontology pre-training (purple, solid). *Plots generated using wandb*.

### Classification Tasks

#### Binary Classification Tasks

For BACE, we see a better performance and convergence on the validation set when using the ontology pre-training (**Fig. 3**). However, different metrics on the test set favour either adding or not adding the ontology pre-training step: while the F1 score is higher without the ontology pre-training step (0.805 without and 0.783 with), the ROC-AUC score is higher with the ontology pre-training (0.872 with and 0.851 without, **Table 5**).

**Table 5.** Test set evaluation for binary classification tasks, averaged over three runs. *Better model in bold*.

| Dataset | OPT | F1 | ROC-AUC |
| --- | --- | --- | --- |
| BACE | w/ | 0.783 $\pm 0.012$ | **0.872** $\pm 0.006$ |
| | w/o | **0.805** $\pm 0.024$ | 0.851 $\pm 0.005$ |
| BBBP | w/ | 0.936 $\pm 0.002$ | 0.955 $\pm 0.001$ |
| | w/o | **0.939** $\pm 0.007$ | **0.962** $\pm 0.008$ |

**Fig. 3.**
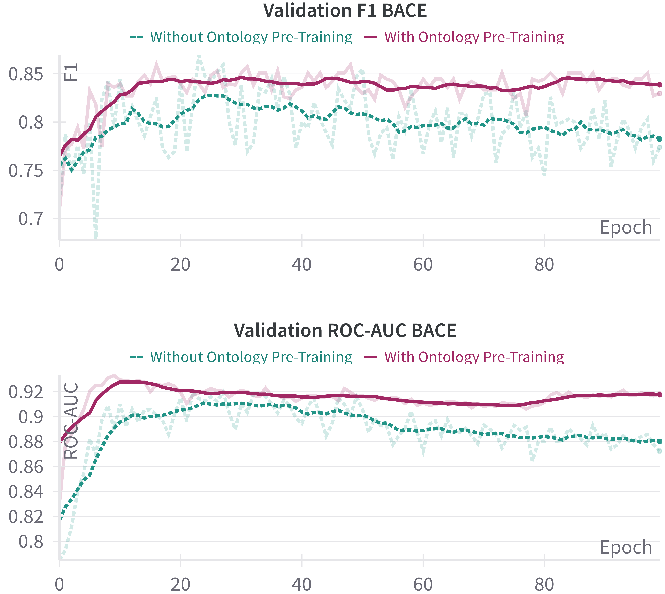
Development of F1, ROC-AUC during training per epoch on the validation dataset for BACE, using a model trained without (turquoise, dashed) and with ontology pre-training (purple, solid). *Plots generated using wandb*.

For BBBP, we see better performance and convergence on the validation set during training when not using the ontology pre-training (**Fig. S4**), however the performance on the test set is better for the model which was not pre-trained on the ontology (**Table 5**).

#### Multi-label Classification Tasks

For SIDER, while we observe a better training performance on the validation set using the ontology pre-training step (**Fig. S5**), the performance on the test set is better for not using the ontology pre-training step (**Table 6**).

**Table 6.** Test set evaluation for multi-label classification tasks, averaged over three runs. *Better model in bold*.

| Dataset | OPT | Micro-F1 | Macro-F1 | ROC-AUC |
| --- | --- | --- | --- | --- |
| SIDER | w/ | 0.601 $\pm 0.004$ | 0.805 $\pm 0.005$ | 0.550 $\pm 0.044$ |
| | w/o | **0.606** $\pm 0.010$ | **0.810** $\pm 0.003$ | **0.612** $\pm 0.031$ |
| ClinTox | w/ | **0.891** $\pm 0.018$ | **0.974** $\pm 0.005$ | **0.991** $\pm 0.002$ |
| | w/o | 0.875 $\pm 0.007$ | 0.971 $\pm 0.002$ | 0.978 $\pm 0.004$ |
| Tox21 | w/ | **0.403** $\pm 0.019$ | **0.463** $\pm 0.012$ | 0.815 $\pm 0.010$ |
| | w/o | 0.370 $\pm 0.021$ | 0.444 $\pm 0.004$ | **0.843** $\pm 0.009$ |

For ClinTox, we see a better convergence, more stable training and better performance when using the ontology pre-training step for the ClinTox dataset (**Table 6, Fig. S6**).

For the Tox21 dataset we again observe higher stability during training as well as delayed and less pronounced over-fitting (**Fig. 4**). Also Micro- and Macro-F1 score based performance on the test set is higher when using the ontology pre-training (**Table 6**). As the Tox21 dataset contains a high amount of missing labels, analogously to our previous work [17], we keep those data points for training but treat any predictions of our model as correct for those missing label data points. This allows us to keep the full dataset for training, without the missing labels affecting our evaluation.

**Fig. 4.**
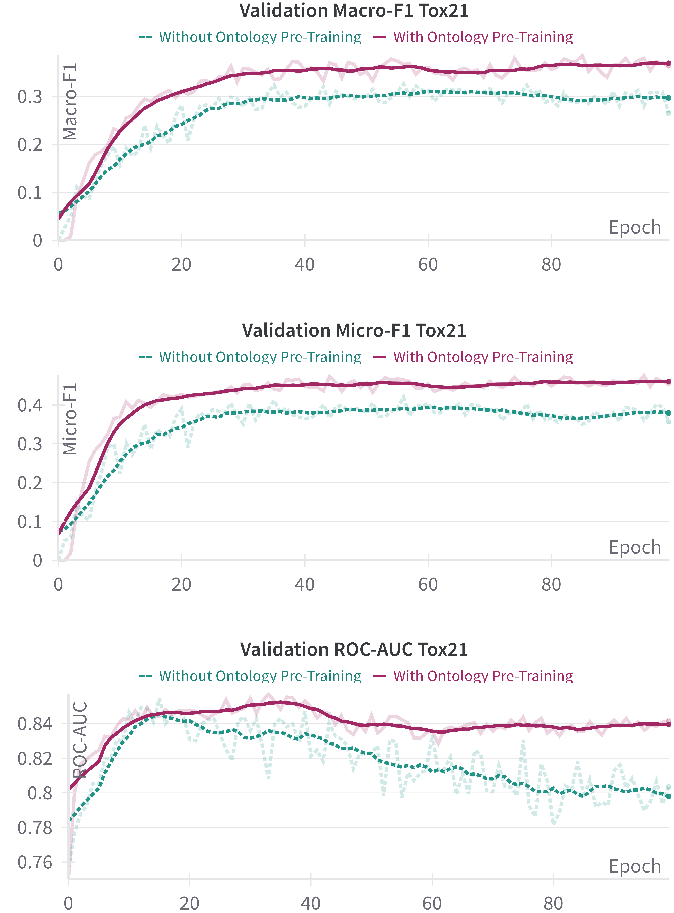
Development of ROC-AUC, Macro-/Micro-F1 during training per epoch on the validation dataset for Tox21, using a model trained without (turquoise, dashed) and with ontology pre-training (purple, solid). *Plots generated using wandb*.

## Discussion

### Comparison to Related Work

#### Non-Machine learning Approaches

We tested our method on a number of downstream prediction tasks with different approaches of experimental determination and applications. For example, aqueous solubility is an important parameter informing bioavailability of e.g. drugs. Some experimental measurements of aqueous solubility are performed using the shake-flask method, column elution or slow-stir method [45, 19]. Various in silico methods have been developed to predict the solubility and a number of other properties of a chemical substance, based on physics-based Quantum Mechanics-Quantitative Structure Property Relationship methods [46]. Similarly, a number of physics-based methods have been used to predict other properties such as lipophilicity using e.g. free energy simulations [47].

#### General Machine Learning Approaches

Many machine learning based methods have been applied to molecular property prediction, using different input types (e.g. SMILES [48], fingerprints [49], molecular graph representation [50]). A number of transformer based models has been suggested [48, 51], as well as graph neural network (GNN) based models [52, 53]. For example, Chemprop [52] uses a graph convolutional model using molecular graphs derived from SMILES as input. Attentive FP uses a graph neural network architecture with a graph attention mechanism [54]. GNNs are currently outperforming other methods, however, struggle to capture long-range interactions.

#### Knowledge Injection Approaches

Knowledge injection can be approached in several different ways, most commonly in the form of feature augmentation based on ontologies [55]. Other approaches include embedding the ontology directly into the network, most commonly by performing a random walk on the ontology [13].

Approaches similar to our ontology pre-training approach have been used in e.g. protein function prediction by performing pre-training using embeddings [56] from the Gene Ontology (GO) [9, 57]. However, while in this method GO is embedded as a knowledge graph to enhance existing protein embeddings, we use ChEBI more directly by using the information in the form of class structure as well as class hierarchy.

Other methods that incorporate expert knowledge into ML-based models to predict properties of molecules have been suggested. KANO [58] integrates functional groups by creating a knowledge graph based on the periodic table and a small number of functional groups from Wikipedia. That graph is then used in the pre-training process by augmenting the molecular graph and in the fine-tuning process by adding prompts to the embeddings.

In [59], it has been shown that integrating functional groups from ChEBI into the KANO model can improve performance for some datasets. Based on the model KANO, a greater number of functional groups are integrated by using information obtained from ChEBI. Two approaches are used to integrate those functional groups: either replacing the original functional group sub-graph from the knowledge graph created by KANO or integrating the functional groups from ChEBI alongside the original ones.

KnoMol adds information about ring systems and topology related information [60], using a graph representation of molecules and knowledge attention heads to incorporate chemical knowledge directly into a transformer model. While it is not using ontology-derived knowledge, the multipersepective attention used includes different attention heads that incorporate expert knowledge about (recognition of) structures such as ring systems and conjugated systems.

We compare our results to the results of different versions of KANO as well as KnoMol. For KANO, we use the results reported in [59] for the original KANO method (KANO,E), the original KANO method without a limit in the group detection (KANO*, E), extended use of functional groups in KANO with the replace approach (KANO*, C), and extended use of functional groups in KANO with the integrate approach (KANO*, E+C). KnoMol only reports test set performance results for the classification problems. In many cases, we outperform those models (**Table 7** for the regression tasks and **Table 8** for the classification tasks). It should be noted that direct comparison are difficult in some cases, due to e.g. the handling of missing labels in the Tox21 dataset (**Table 2**). In addition, the only reported metric is ROC-AUC, which is misleading for imbalanced datasets [42]: a comparison using metrics such as micro- and macro-F1 score would be more appropriate.

**Table 7.** RMSE of regression problems, as reported by other methods. Better models in bold. OPT-ChEB-AI: Ontology Pre-Training-ChEB-AI.

| Method | ESOL | FreeSolv | Lipophilicity |
| --- | --- | --- | --- |
| OPT-ChEB-AI | **0.719** | **1.367** | 0.750 |
| KANO, E [58, 59] | 0.776 | 1.593 | 0.610 |
| KANO*, E [59] | 0.766 | 1.595 | **0.600** |
| KANO*, C [59] | 0.747 | 1.675 | 0.612 |
| KANO*, E+C [59] | 0.740 | 1.584 | 0.601 |
| KnoMol [60] | n/a | n/a | n/a |

**Table 8.**
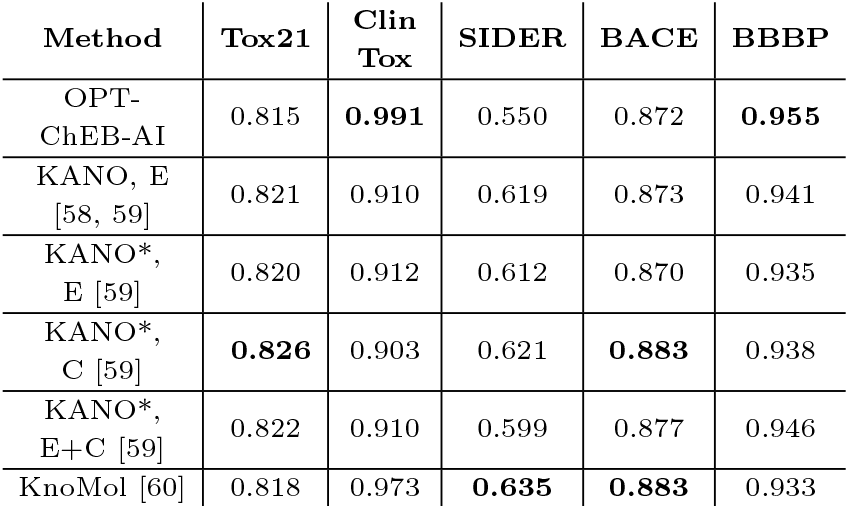
ROC-AUC of classification problems, as reported by other methods. Better models in bold. OPT-ChEB-AI: Ontology Pre-Training-ChEB-AI.

| Method | Tox21 | Clin<br>Tox | SIDER | BACE | BBBP |
| --- | --- | --- | --- | --- | --- |
| OPT-ChEB-AI | 0.815 | **0.991** | 0.550 | 0.872 | **0.955** |
| KANO, E<br>[58, 59] | 0.821 | 0.910 | 0.619 | 0.873 | 0.941 |
| KANO*,<br>E [59] | 0.820 | 0.912 | 0.612 | 0.870 | 0.935 |
| KANO*,<br>C [59] | **0.826** | 0.903 | 0.621 | **0.883** | 0.938 |
| KANO*,<br>E+C [59] | 0.822 | 0.910 | 0.599 | 0.877 | 0.946 |
| KnoMol [60] | 0.818 | 0.973 | **0.635** | **0.883** | 0.933 |

### Benefit of Ontology Pre-Training

Adding an ontology pre-training step leads to more robust models. For 6 out of 9 datasets, adding the ontology pre-training step improves performance, for the remaining datasets the performance is mostly similar with and without the ontology pre-training step. For 4 of 8 datasets, our method achieves better performance than other methods. While the results are not beating or reaching the state-of-art models for every dataset, we were able to show the benefit of adding an ontology pre-training step into the pipeline. In many cases we see better convergence, more stable training, and better performance. Especially for the regression problems, adding an ontology pre-training step improved the results every time, also for more difficult classification tasks the results seem to be improved by the knowledge-injection. Our hypothesis is therefore that the harder the problem is the more the ontology pre-training helps.

Furthermore, we hypothesise that our method improves performance for those applications that are in one way or another contained in ChEBI. For example, toxicity and solubility are properties with either classes directly present in ChEBI or associated with certain branches of the ontology. Therefore, it would be interesting to investigate the effect of using more directly related ontologies for those predictive tasks where we did not see an improvement with the ontology pre-training.

Additionally, our method is more generalisable than others, as ontologies crafted by experts can be easily incorporated into the training process, only requiring that the underlying input data type for the model can be associated with the ontology classes in some way. This also allows for updates in the ontology to be easily transferred to the model by re-training the pre-trained model on the updated ontology. Our turnkey usage of ontologies improves down-stream applications without the need for feature augmentation such as incorporating handcrafted features.

### Limitations

As we already observed in our previous work [17], deep-learning models which are trained on small biological datasets often suffer from over-fitting. Furthermore, we do not always see an improvement for all the classification problems we explored.

It is, however, well known that the ROC-AUC metric does not perform well on unbalanced datasets [42]; in our study the Tox21 and BBBP datasets in particular have a strong imbalance in terms of target label distribution (**Table 2**). In addition, all possible assignments of the prediction threshold are considered when calculating the ROC-AUC value. However, this threshold is a crucial hyperparameter for the prediction quality of the model. These parameters should rather be selected based on the training and validation data, as adjusting them to the test data negatively affects the significance of the results. Therefore, we consider thresholded metrics such as *F*_*β*_ - metrics more meaningful when evaluating the final predictive performance of a model, while ROC-AUC is an indicator for model sensitivity.

### Future Work

We would like to extend our methods to further applications, such as property prediction for proteins or RNA, injecting knowledge from the Gene Ontology [9] or RNA-KG [61] similarly to our approach here. Furthermore, as discussed above, adding an ontology pre-training step to state-of-the-art models might greatly improve overall performance. In addition, more advanced architectures such as Graph Neural Networks may be capable to further exploit the hierarchical structure of ontologies, leading to improvements in ontology-derived knowledge injection. Finally, we aim to improve ring system representation, similarly to methods such as presented in [60].

## Conclusion

In [17], we have shown that adding an ontology pre-training step has benefits for solving the problem of classification of toxicity of molecules. Here, we have extended this method to regression problems as well as other classification problems. Hence, we contribute to establishing ontology pre-training as a widely applicable method for improving the performance of machine learning models. We outperform similar knowledge-injection methods for 4 of 8 MoleculeNet datasets [18]. Adding an ontology pre-training step improves training stability and performance in 6 of 9 of our datasets. Our intuitive and extensible method therefore shows promise for many additional applications.

## Acknowledgments

This work was supported by the Swiss National Science Foundation, grant agreement number 215906 and by the Deutsche Forschungsgesellschaft (DFG, German Research Foundation) - 522907718.

## Footnotes

1 https://github.com/ChEB-AI/python-chebai

2 https://doi.org/10.5281/zenodo.17202668

