## Supplementary Material for "Ontology pre-training improves machine learning-based predictions for metabolites"

### Plots with validation set metrics during training

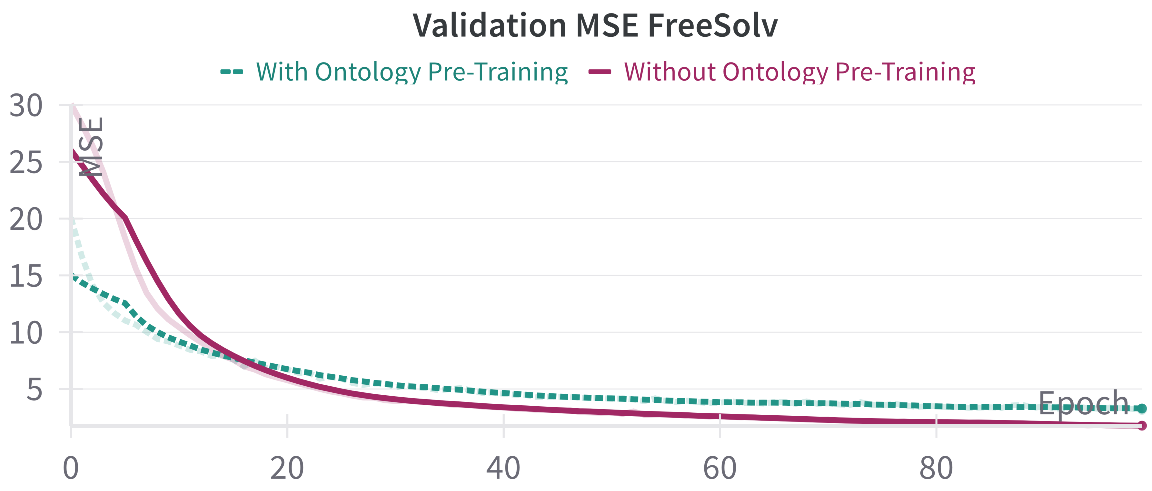


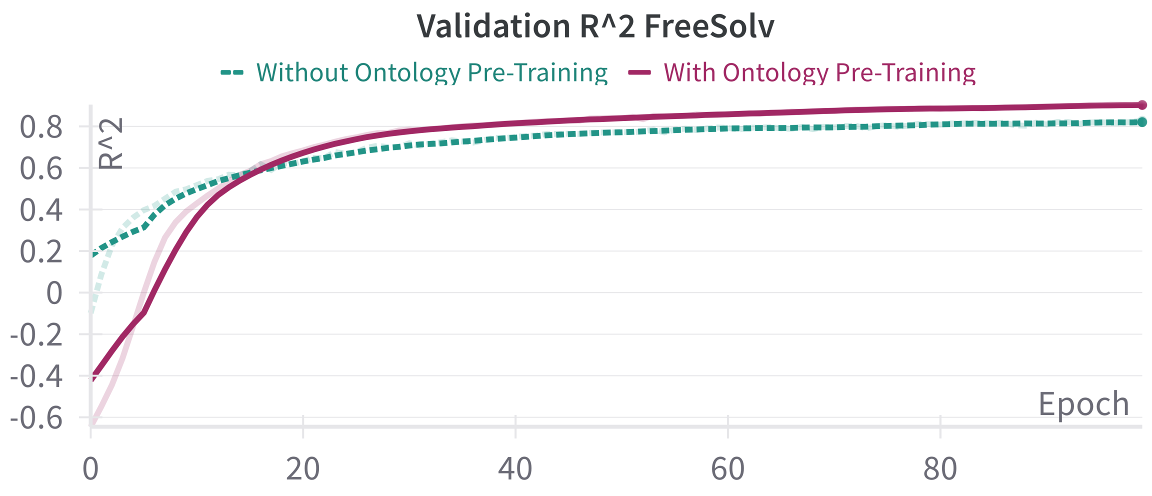


**Supplementary Figure 1.** Development of MSE, $R^2$ during training per epoch on the validation dataset for FreeSolv, using a model trained without (turquoise, dashed) and with ontology pre-training (purple, solid). *Plots generated using wandb.*

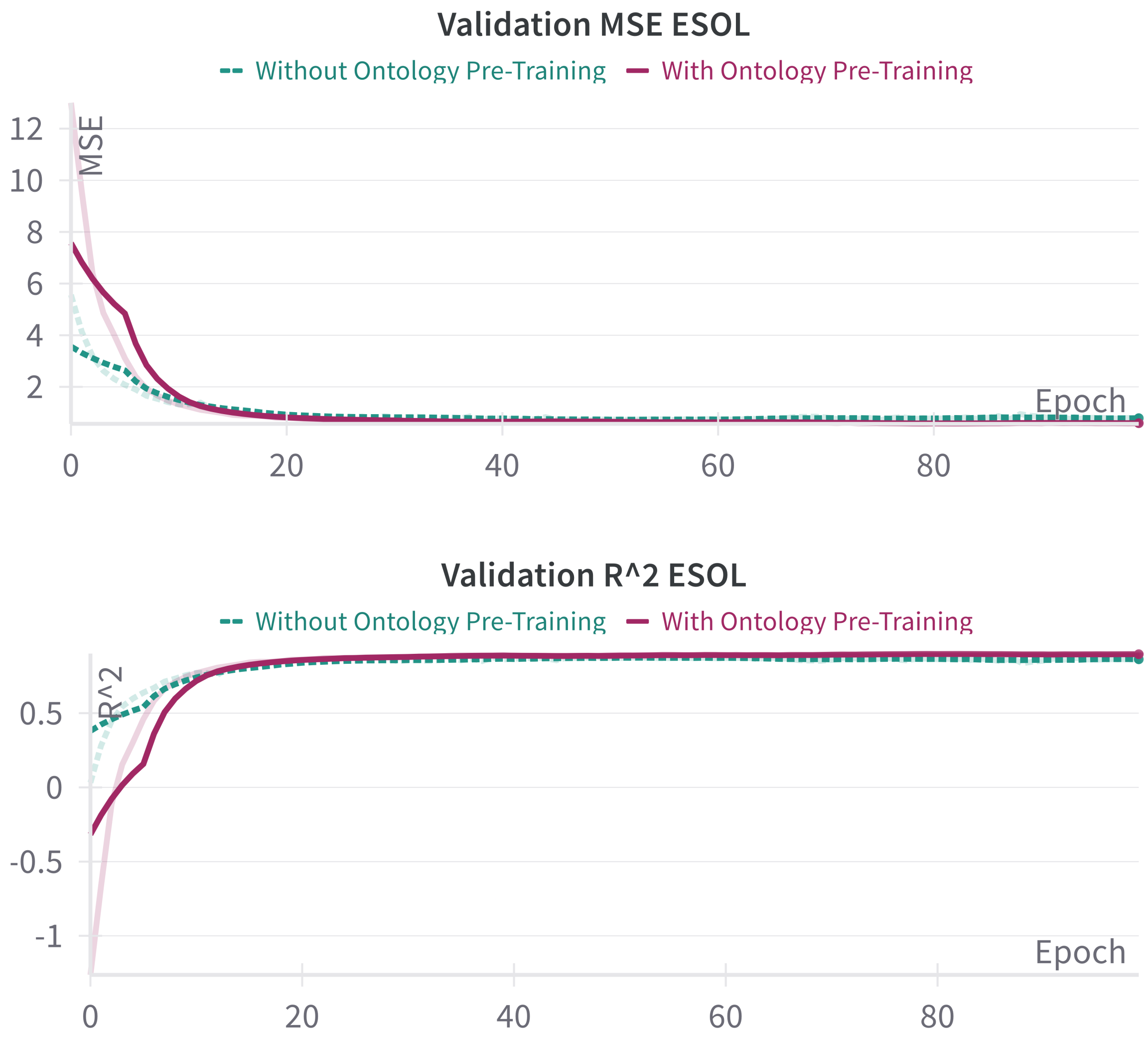


**Supplementary Figure 2.** Development of MSE, $R^2$ during training per epoch on the validation dataset for ESOL, using a model trained without (turquoise, dashed) and with ontology pre-training (purple, solid). *Plots generated using wandb.*

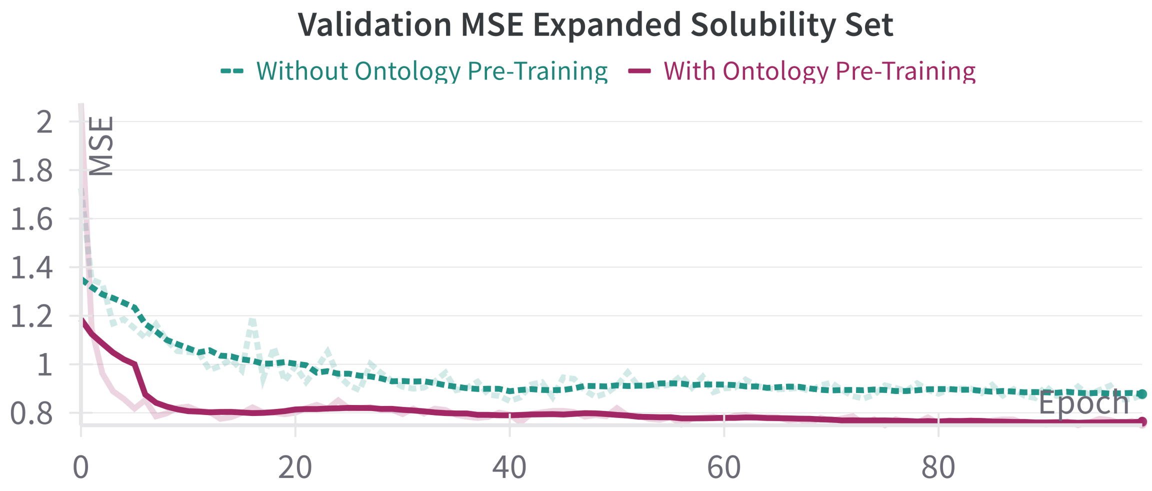


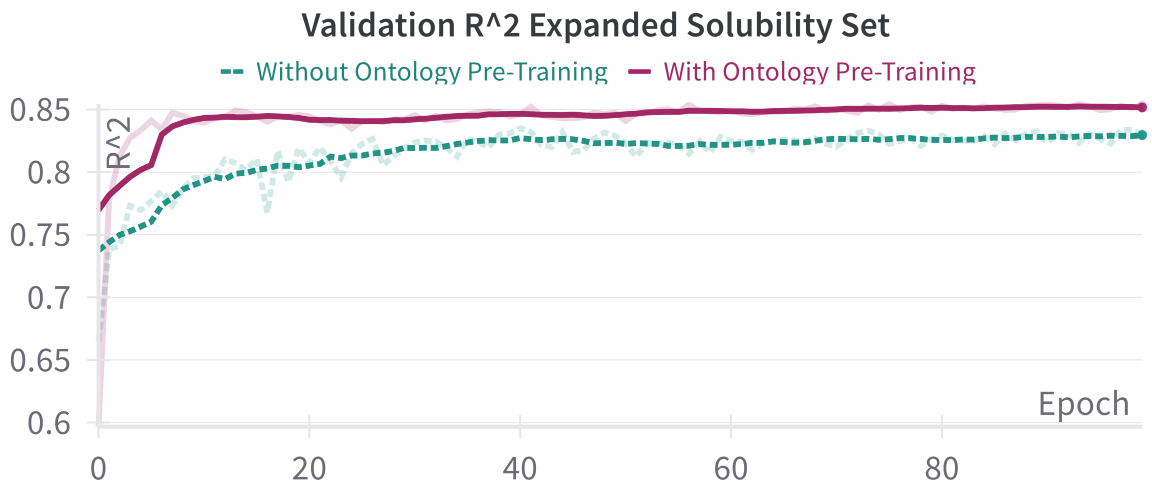


**Supplementary Figure 3.** Development of MSE, $R^2$ during training per epoch on the validation dataset for Solubility, using a model trained without (turquoise, dashed) and with ontology pre-training (purple, solid). *Plots generated using wandb.*

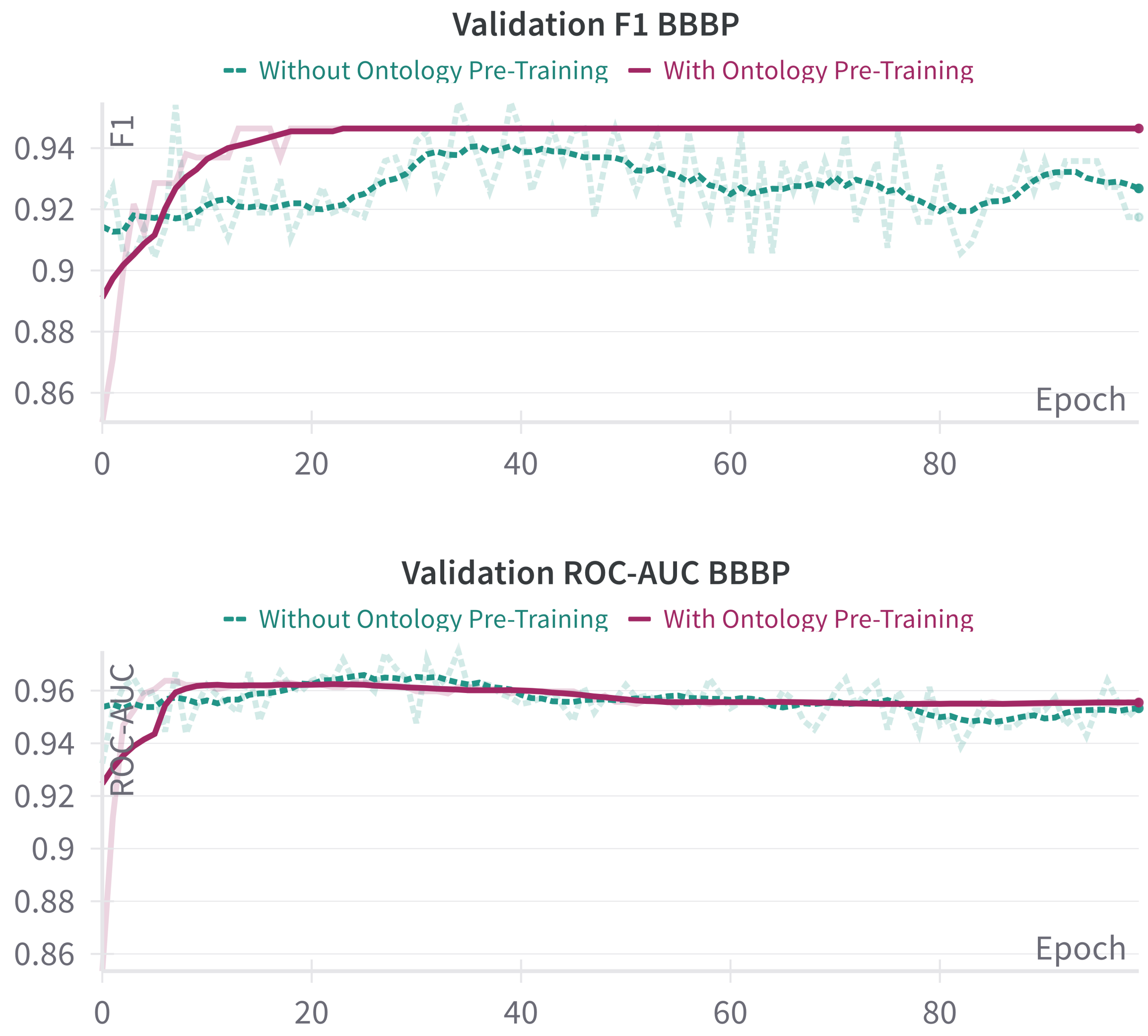


**Supplementary Figure 4.** Development of F1, ROC-AUC during training per epoch on the validation dataset for BBBP, using a model trained without (turquoise, dashed) and with ontology pre-training (purple, solid). *Plots generated using wandb.*

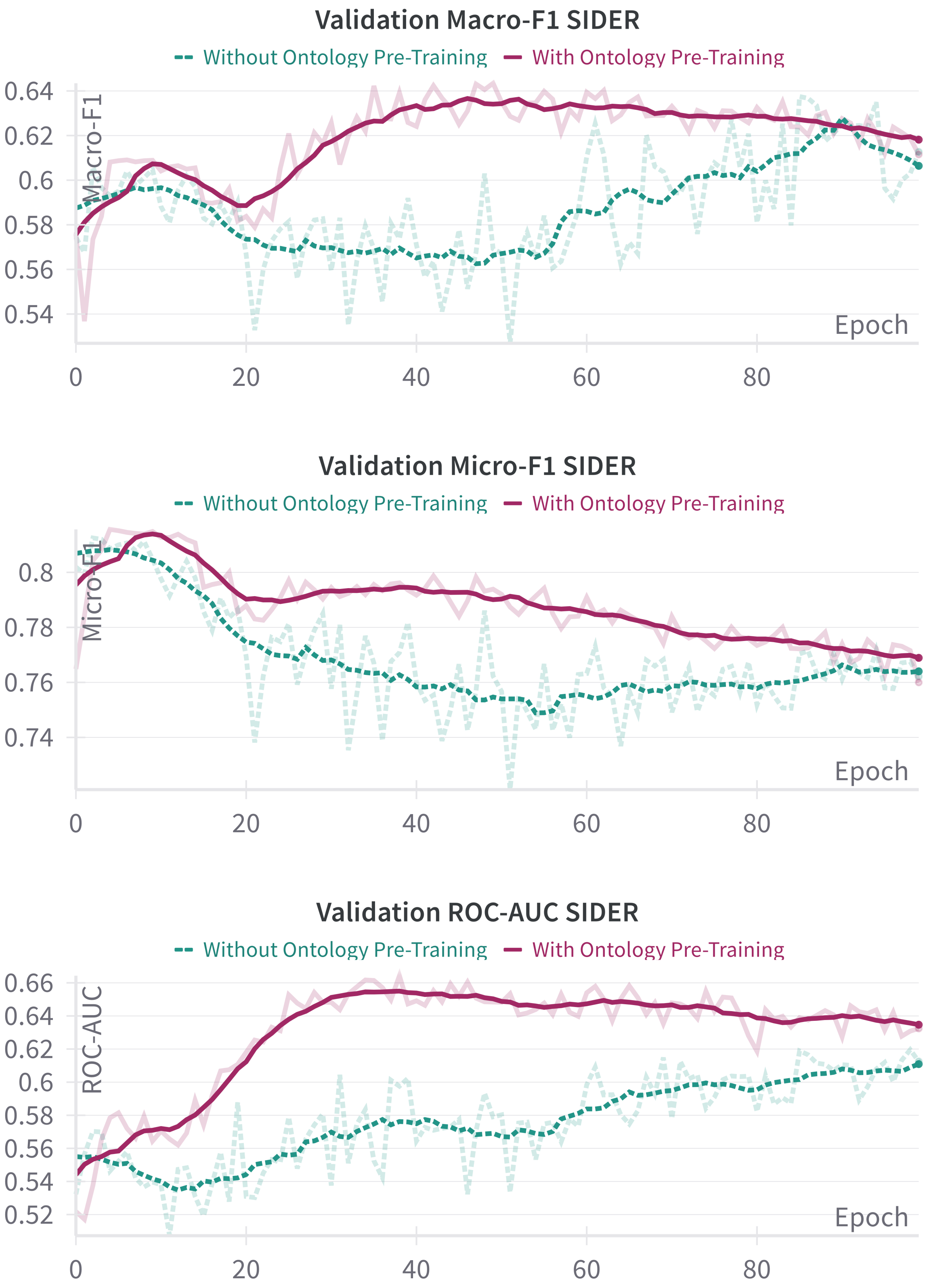


**Supplementary Figure 5.** Development of (macro-/micro-)F1, ROC-AUC during training per epoch on the validation dataset for SIDER, using a model trained without (turquoise, dashed) and with ontology pre-training (purple, solid). *Plots generated using wandb.*

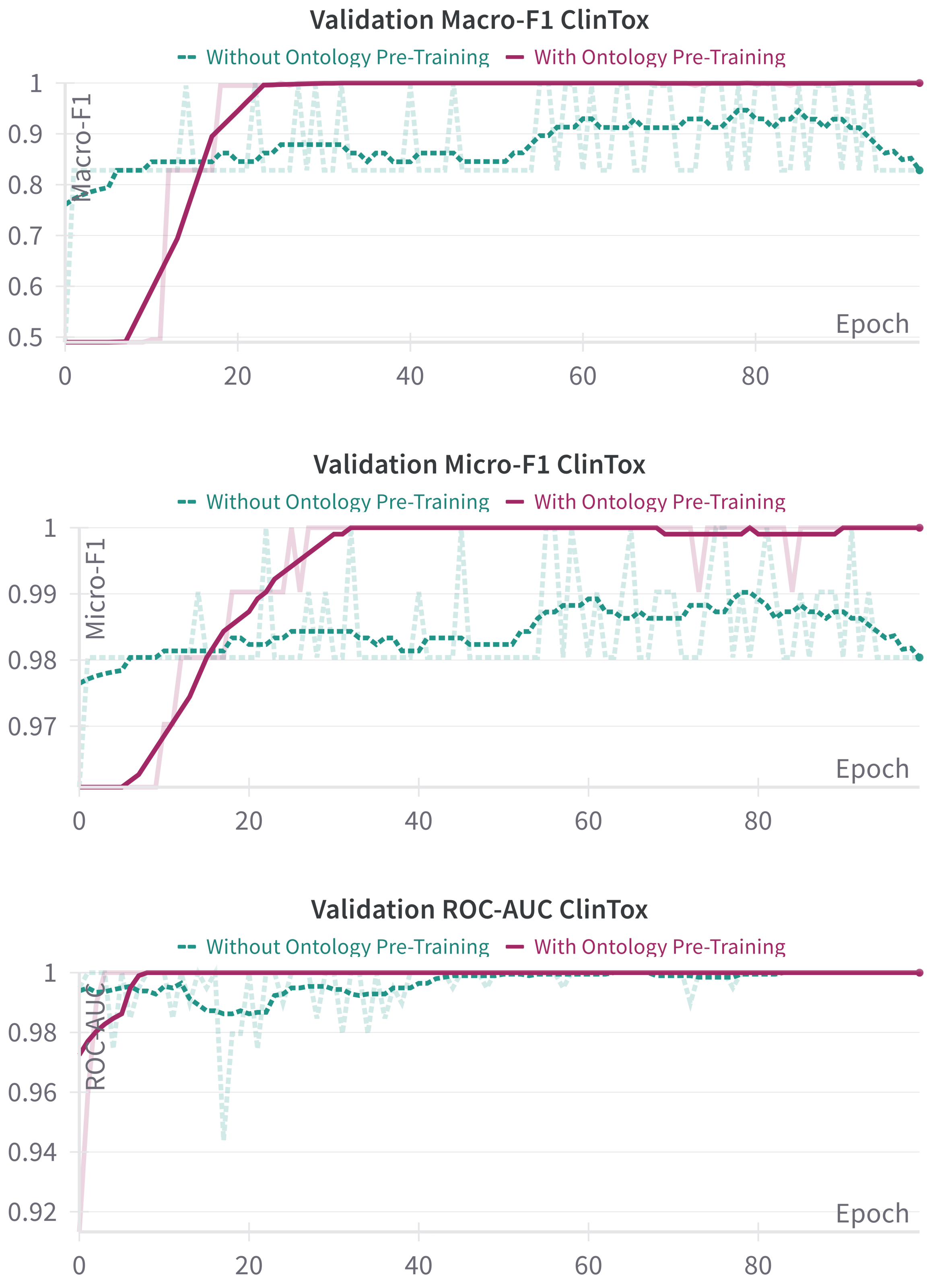


**Supplementary Figure 6.** Development of (macro-/micro-)F1, ROC-AUC during training per epoch on the validation dataset for ClinTox, using a model trained without (turquoise, dashed) and with ontology pre-training (purple, solid). *Plots generated using wandb.*

### Datasets

| Dataset | Description | Type | Task (#Labels) | #Compounds |
|---|---|---|---|---|
| Lipophlicity | Membrane permeability and solubility | Physical chemistry | Regression | 4,200 |
| FreeSolv | Hydration free energy of small molecules in water | Physical chemistry | Regression | 643 |
| ESOL | Aqueous solubility | Physical chemistry | Regression | 1,128 |
| Expanded Solubility dataset | Several curated solubility datasets | Physical chemistry | Regression | 17,339 |
| BACE | Binding results for a set of inhibitors | Biophysics | Classification (1) | 1,522 |
| BBBP | Blood-brain barrier permeability property | Physiology | Classification (1) | 2,053 |
| SIDER | Adverse drug reactions | Physiology | Classification (27) | 1,427 |
| ClinTox | Drug's toxicity and FDA approval | Physiology | Classification (2) | 1,491 |
| Tox21 | Toxicity | Physiology | Classification (12) | 8,014 |

**Table 1.** Overview of datasets used for fine-tuning.

### Results for more metrics

| Dataset | OPT | RMSE | $R^2$ |
|---|---|---|---|
| FreeSolv | w/ | **1.367** | **0.880** |
| | w/o | 1.570 | 0.841 |
| Lipophilicity | w/ | **0.750** | **0.591** |
| | w/o | 0.905 | 0.403 |
| ESOL | w/ | **0.719** | **0.863** |
| | w/o | 0.832 | 0.817 |
| Expanded Solubility Dataset | w/ | **0.869** | **0.859** |
| | w/o | 0.978 | 0.821 |

**Table 2.** Test set evaluation for regression tasks. $R^2$ and RMSE metrics, averaged over three runs. Better model in bold.

| Dataset | OPT | F1 | ROC-AUC | PRC-AUC | Bal. Acc. |
|---|---|---|---|---|---|
| BACE | w/ | 0.783 | **0.872** | **0.847** | 0.783 |
| | w/o | **0.805** | 0.851 | 0.817 | **0.797** |
| BBBP | w/ | 0.936 | 0.955 | 0.987 | 0.851 |
| | w/o | **0.939** | **0.962** | **0.990** | **0.897** |

**Table 3.** Test set evaluation for binary classification tasks, averaged over three runs. Better model in bold.

| Dataset | OPT | Micro-F1 | Macro-F1 | ROC-AUC | PRC-AUC | Bal. Acc. |
|---|---|---|---|---|---|---|
| SIDER | w/ | 0.601 | 0.805 | 0.550 | 0.619 | 0.503 |
| | w/o | **0.606** | **0.810** | **0.612** | **0.646** | **0.511** |
| ClinTox | w/ | **0.891** | **0.974** | **0.991** | **0.966** | 0.836 |
| | w/o | 0.875 | 0.971 | 0.978 | 0.941 | **0.874** |
| Tox21 | w/ | **0.403** | **0.463** | 0.815 | **0.394** | **0.666** |
| | w/o | 0.370 | 0.444 | **0.843** | 0.393 | 0.651 |

**Table 4.** Test set evaluation for multi-label classification tasks, averaged over three runs. Better model in bold.
